# The Flowering Time Regulator *FLK* Controls Pathogen Defense in *Arabidopsis thaliana*

**DOI:** 10.1101/2021.01.10.426133

**Authors:** Matthew Fabian, Min Gao, Xiao-Ning Zhang, Jiangli Shi, Sung-Ha Kim, Priyank Patel, Anna R. Hu, Hua Lu

**Affiliations:** Department of Biological Sciences, University of Maryland Baltimore County, 1000 Hilltop Circle, Baltimore, MD 21250, USA; Biochemistry Program, Department of Biology, St Bonaventure University, St Bonaventure, NY 14778, USA; Department of Biology Education, Korea National Univ. of Education, Chungbuk, KOREA

## Abstract

Plant disease resistance is a complex process that is maintained in an intricate balance with development. Increasing evidence indicates the importance of post-transcriptional regulation of plant defense by RNA binding proteins. The K homology (KH) repeat is an ancient RNA binding motif found in proteins from diverse organisms. The role of KH domain proteins in pathogen resistance is not well known. From a genetic screen aimed to uncover novel defense genes in Arabidopsis, we identified a new allele of the canonical flowering regulatory gene, *FLOWERING LOCUS KH Domain* (*FLK*), encoding a putative triple KH-repeat protein. In addition to late flowering, the *flk* mutants exhibited decreased resistance to the bacterial pathogen *Pseudomonas syringae* and increased resistance to the necrotrophic fungal pathogen *Botrytis cinerea*. We found that the *flk* mutations compromised basal defense and defense signaling mediated by salicylic acid and led to increased reactive oxygen species (ROS) scavenging, likely through *FLK*’s regulation of the ROS scavenging enzyme catalases. RNA-seq data revealed that major defense signaling genes are regulated by *FLK*, providing a molecular basis for *FLK*’s contribution to pathogen defense. Together our data support that FLK is a multifunctional protein regulating pathogen defense and development of plants.

## Introduction

Plants are constantly challenged by pathogens with different lifestyles. Plants can recognize pathogen-derived molecules, such as pathogen associated molecular patterns (PAMPs) and effector proteins, and subsequently activate layers of defense responses, including PAMP-triggered immunity (PTI), effector-triggered immunity, and systemic acquired resistance (Boller and Felix, 2009; Dodds and Rathjen, 2010; Fu and Dong, 2013; Toruno et al., 2016). Depending on the type of invading pathogen, plants deploy different and sometimes conflicting downstream signaling to combat the invader. For instance, salicylic acid (SA) is generally considered to be important for defense against biotrophic pathogens, e.g., *Pseudomonas syringae*, whereas jasmonic acid (JA) promotes resistance to necrotrophic pathogens, e.g., *Botrytis cinerea*, as well as insects (Glazebrook, 2005; Campos et al., 2014). SA and JA act antagonistically in the defense response against some pathogens, but they also work synergistically under other stress conditions. SA is also known to crosstalk with reactive oxygen species (ROS) (Herrera-Vásquez et al., 2015). However, the role of ROS in pathogen defense is less well understood. Some studies suggest that reduced ROS is associated with *P. syringae* susceptibility and *Botrytis* resistance in plants (Chamnongpol et al., 1998; Govrin and Levine, 2000; Polidoros et al., 2001; Rossi et al., 2017; Yuan et al., 2017); but these observations are challenged by some other studies (Macho et al., 2012; Li et al., 2014; Survila et al., 2016). Proper interplays among signaling pathways are important to ensure robust plant defense. Without a thorough understanding of how different signaling molecules activate defense against pathogens, our design of strategies in improving disease resistance of crop plants against their natural pathogens will be limited.

Defense is an energetically costly process and defense activation against some pathogens can come at the expense of plant development (Karasov et al., 2017; Ning et al., 2017). Flowering is one of the most critical developmental landmarks in the lifecycle of plants. Arabidopsis flowering is tightly controlled by at least five pathways: the autonomous pathway, photoperiod (light), vernalization, hormones, and the circadian clock (Amasino, 2010). These pathways ultimately converge upon a few downstream genes, e.g., the flowering repressor gene *FLOWERING LOCUS C* (*FLC*) and activator genes *FT* and *SUPPRESSOR OF OVEREXPRESSION OF CONSTANS1*, to determine flowering time. Growing evidence supports the involvement of flowering pathways in defense, including the autonomous pathway genes *FPA* and *FLD* (Lyons et al., 2013; Singh et al., 2013); light and certain light receptors (Genoud et al., 2002; Griebel and Zeier, 2008; Roden and Ingle, 2009); flowering regulatory hormones (Spoel and Dong, 2008; Bari and Jones, 2009); and the circadian clock (Lu et al., 2017). On the other hand, defense activation reciprocally affects flowering. For instance, pathogen infection (Korves and Bergelson, 2003) and changes in SA levels via exogenous SA application or genetic manipulation are associated with altered flowering time (Martinez et al., 2004; Jin et al., 2008; March-Diaz et al., 2008; Endo et al., 2009; Wada et al., 2009; Liu et al., 2012). These observations suggest crosstalk between plant defense and flowering control. Mechanisms underlying this crosstalk remain to be fully elucidated.

*FLOWERING LOCUS K Homology Domain* (*FLK*) is an autonomous pathway flowering gene encoding a protein with three K homology (KH) repeats (Lim et al., 2004; Mockler et al., 2004). The KH domain is an ancient RNA binding motif found in proteins from diverse organisms (Nicastro et al., 2015). KH domain proteins have up to 15 KH repeats and some are known to function in RNA metabolism, such as pre-mRNA processing and mRNA stability. Disruption of KH domain proteins is associated with multiple diseases in humans (Geuens et al., 2016). There are 26 KH domain proteins in Arabidopsis (Lorkovic and Barta, 2002) but only a few of them have been characterized. Like FLK, several KH domain proteins were shown to play roles in flowering control (Cheng et al., 2003; Ripoll et al., 2009; Yan et al., 2017; Ortuno-Miquel et al., 2019; Woloszynska et al., 2019). Some KH domain proteins were shown to be important for stress response, including resistance against a fungal pathogen (Thatcher et al., 2015) and abiotic stress (Chen et al., 2013; Guan et al., 2013). Mechanisms underlying the functions of these KH proteins remain largely unknown. To date, none of the KH domain proteins are shown to influence both defense and development in Arabidopsis.

The Arabidopsis mutant *acd6-1* exhibits a constitutively high level of defense that is inversely correlated with the size of the plant (Rate et al., 1999; Lu et al., 2003). This feature of *acd6-1* makes it a powerful genetic tool in uncovering novel defense genes in a large-scale genetic screen for *acd6-1* suppressors (Lu et al., 2009; Wang et al., 2014). *acd6-1* has also been conveniently used to gauge interactions among defense genes in genetic analyses (Song et al., 2004; Ng et al., 2011; Wang et al., 2011a; Wang et al., 2014; Hamdoun et al., 2016; Zhang et al., 2019). From the genetic screen, we isolated a new allele of the *FLK* gene, *flk-5*, that suppressed *acd6-1-*conferred phenotypes, including SA accumulation, cell death, and dwarfism. In addition to late flowering, *flk* mutants showed enhanced susceptibility to the biotrophic pathogen *P. syringae* but enhanced resistance to the necrotrophic pathogen *Botrytis*. In addition, the *flk* mutants exhibited compromised SA accumulation and basal defense upon *P. syringae* infection and/or elicitation by *P. syringae* derived molecules. The *flk* mutants further displayed reduced ROS accumulation, likely due to an increased ROS scavenging. RNA-seq analysis supported the roles of *FLK* in regulating pathogen defense and development. Together our data establish that *FLK* is a multifunctional gene important for plant defense and development.

## Results

### Identification of a new *FLK* allele from *acd6-1* suppressor screen

From a large-scale mutant screen for *acd6-1* suppressors generated by T-DNA insertional mutagenesis, we isolated a new allele of the *FLK* gene that has a T-DNA insertion in the second intron of the gene (Figure 1A). *FLK* encodes a protein with three KH repeats and is known as an autonomous pathway gene regulating flowering (Lim et al., 2004; Mockler et al., 2004). Because four other *flk* mutants were previously described (Lim et al., 2004; Mockler et al., 2004), we designated this mutant as *flk-5*. Compared with *acd6-1, acd6-1flk-5* was larger, had reduced cell death and SA accumulation, and flowered later (Figure 1B-1D). These phenotypes of *acd6-1flk-5* were confirmed by introducing another *FLK* allele, *flk-1* (SALK_007750), into *acd6-1*. In addition, a construct carrying the 2.3 kb *FLK* promoter and the full length *FLK* genomic fragment that was translationally fused with the reporter gene *GFP* (*FLK-GFP*) rescued delayed flowering in *flk-1* (Supplemental Figure 1A and 1B). The FLK*-*GFP protein was localized to the nucleus (Supplemental Figure 1C).

**Figure 1.**
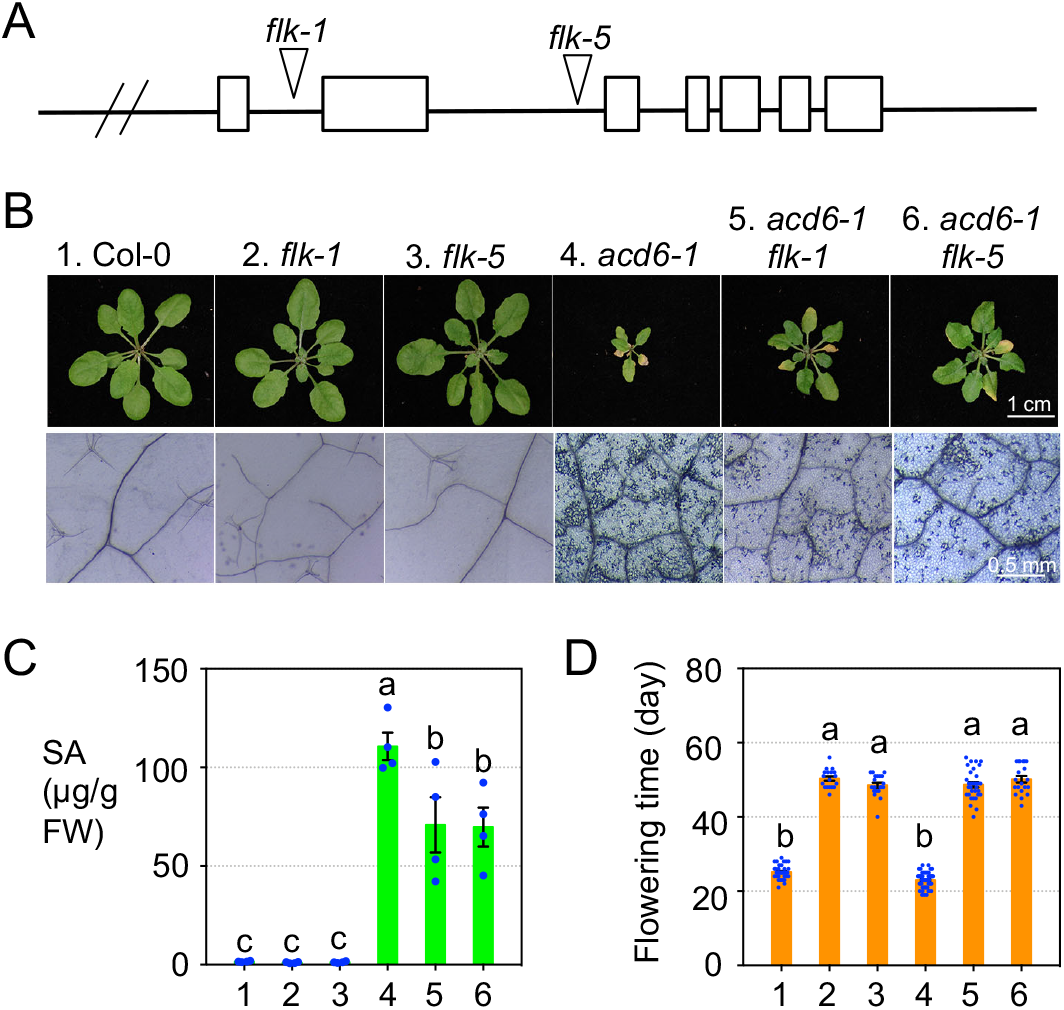
Mutations in *FLK* suppress *acd6-1*-conferred phenotypes and lead to late flowering. (A) Gene structure of *FLK*. Triangles indicate the positions of two *FLK* alleles. (B) Pictures of 25-d old plants (top) and cell death (bottom). Leaves at the fourth to seventh positions of each genotype were stained with trypan blue for cell death. (C) SA quantification. Total SA was extracted from the plants and quantified by HPLC. (D) Flowering time measurement. Plants grown in a light cycle of 16 h L/8 h D were recorded for flowering time, days after planting for the first visible appearance of inflorescence of a plant. Error bars represent mean ± SEM in (C) (n=4) and (D) (n≥18). Different letters in (C) and (D) indicate significant difference among the samples (P<0.05; One-way ANOVA with post-hoc Tukey HSD test). These experiments were repeated two times with similar results.

### *FLK* is a positive regulator of *P. syringae* resistance and a negative regulator of *Botrytis* resistance

In response to invading pathogens of varying lifestyles, plants activate downstream defense signaling that can sometimes act antagonistically. SA and JA are two important defense signaling molecules that are known to play opposing roles in plant defense against some biotrophic and necrotrophic pathogens (Glazebrook, 2005; Campos et al., 2014). The suppression of *acd6-1-*conferred phenotypes by *flk* mutants supports an SA regulatory role for *FLK* and suggests that *FLK* is important for defense against *P. syringae*. To test this, we infected Col-0, *flk-1*, and *flk-5* plants with the virulent *P. syringae* pv. *maculicola* ES4326 strain DG3 (*Pma*DG3). We found that both *flk* mutants were more susceptible than Col-0 and this susceptibility was rescued by *FLK-GFP*, as demonstrated in two representative rescued lines #7 and #20 (Figure 2A). The *flk* mutants also had lower SA accumulation and lower expression of the SA marker gene *PR1* than Col-0 with *Pma*DG3 infection (Figure 2B-2C), further supporting the SA regulatory role of *FLK*.

**Figure 2.**
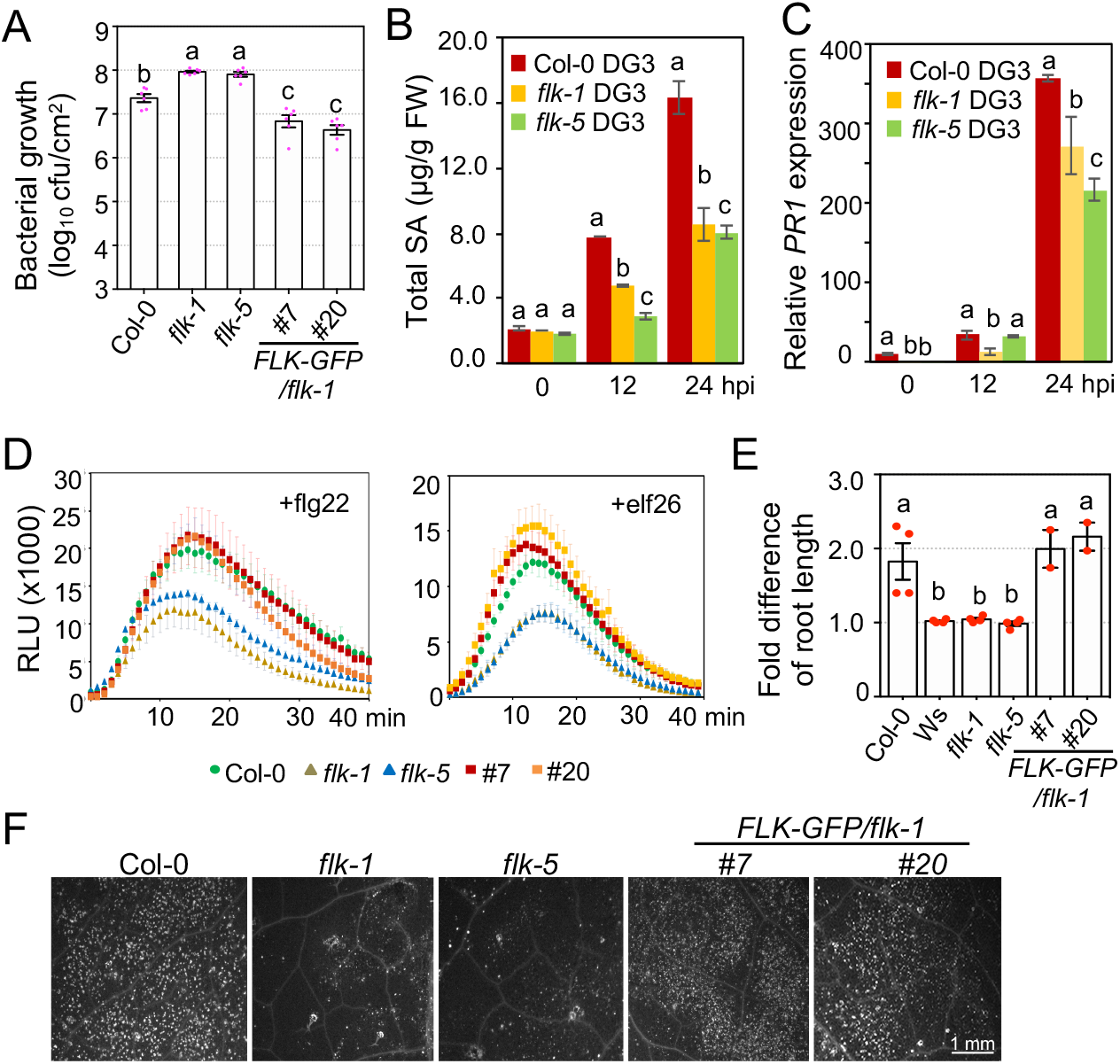
*FLK* positively regulates *P. syringae* resistance, SA accumulation, and PTI. (A) Bacterial growth with *PmaDG3* infection (n=6). (B) SA quantification with *PmaDG3* infection (n=2). (C) Expression of *PR1* with *PmaDG3* infection (n=3). (D) ROS burst in seedlings treated with 1 µM flg22 (left) and 1 µM elf26 (right) (n=12). RLU: relative luminescence unit. (E) Root growth inhibition with 1 µM flg22 treatment. 4-d old seedlings were treated with 1 µM flg22 and measured for root length 4 d post treatment. The flg22-insensitive ecotype Ws that contains a loss-of-function mutation in the *FLS2* gene was included as a control. The fold difference is the ratio of water-treated and flg22-treated root length of each genotype. Data represent the average of three independent experiments. (F) Images of callose deposition. The fourth to seventh leaves of 25-d old plants were infiltrated with 1µM flg22 for 24 h followed by callose staining, using 0.01% aniline blue. Error bars represent mean ± SEM in (A), (D), and (E) and mean ± STDV in panel (B) and (C). Different letters indicate significant difference among the samples of the same time point (P<0.05; One-way ANOVA with post-hoc Tukey HSD test). These experiments were repeated two times with similar results.

Plant perception of biotrophic pathogens is often associated with activation of PTI, a basal level of defense (Zipfel et al., 2004). Flg22, a 22-aa peptide from the conserved region of the flagellin protein of *P. syringae*, is widely used to elicit PTI. Within minutes of recognition of flg22, plants activate an oxidative burst, producing a high level of ROS. We found that the *flk* mutations compromised the ROS burst, compared with Col-0 and the two rescued lines (Figure 2D, left). This phenotype was further corroborated with another PAMP molecule, elf26, a 26-aa peptide from the elongation factor Tu protein (Kunze et al., 2004) (Figure 2D, right). PTI responses also include cell wall strengthening via callose deposition and reduced plant growth. We found that the *flk* mutants displayed fewer callose depositions and less seedling growth inhibition upon flg22 treatment (Figures 2E and 2F). These *flk-*conferred PTI phenotypes were rescued by *FLK-GFP* (Figures 2D-2F). Together our data suggest that *FLK* positively regulates *P. syringae* resistance through affecting SA signaling and PTI.

Interestingly, we found that the *flk* mutants were more resistant to the necrotrophic fungal pathogen *Botrytis cinerea*, compared with Col-0 and the two rescued lines (Figure 3A and 3B). Although JA signaling is known to be involved in *Botrytis* resistance, the *flk* mutants showed a similar response to JA-induced root inhibition (Figure 3C). Therefore, we conclude that *FLK* negatively regulates *Botrytis* resistance likely independently of JA signaling.

**Figure 3.**
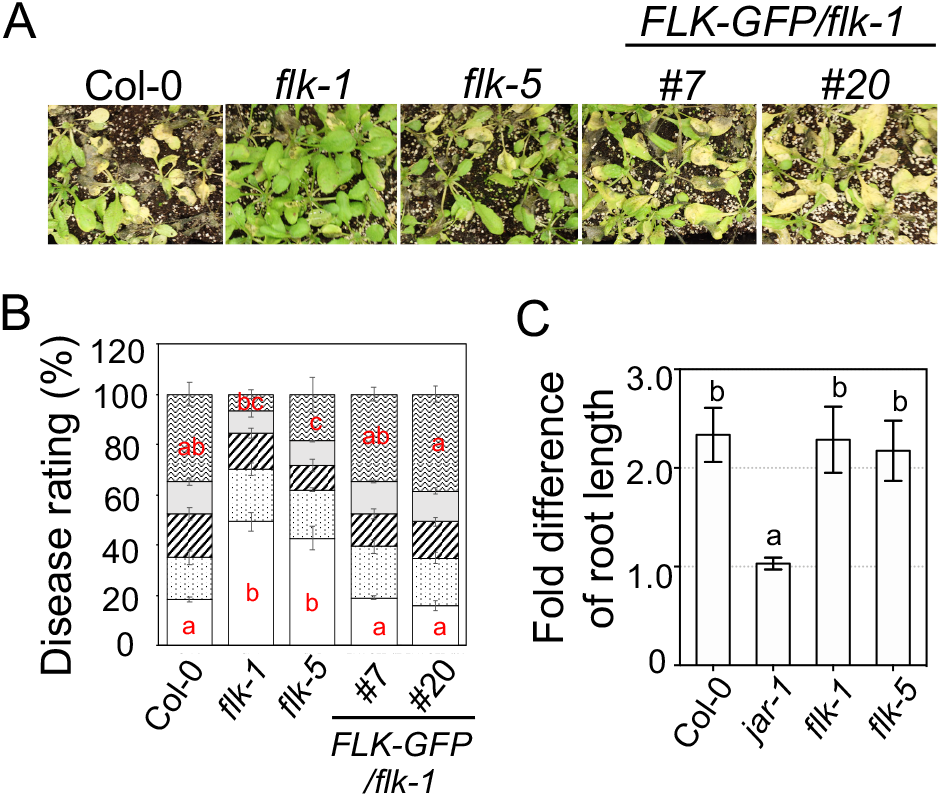
*FLK* negatively regulates *Botrytis* resistance independently of JA signaling. (A) Pictures of *Botrytis-*infected plants. (B) Disease symptom scoring with *Botrytis* infection. The rating scale is from 1 (bottom; no lesion or small, rare lesions) to 5 (top; lesions on over 70% of a leaf). (C) Root inhibition assay. Seedlings were treated with 10 µM methyl jasmonate (MJ). The MJ insensitive mutant *jar-1* was used as a control. Error bars represent mean ± SEM in (B) (n=30) and (C) (n=12). Different letters indicate significant difference among the samples in the same comparison group (P<0.05; One-way ANOVA with post-hoc Tukey HSD test). These experiments were repeated two times with similar results.

### FLK interacts with multiple SA regulators in affecting *acd6-1-*conferred phenotypes

A number of genes are known to contribute to SA-mediated defense. These genes can be grossly grouped into three types based on their functions (Lu, 2009). The type I genes, e.g., *ICS1* and *WIN3* (Wildermuth et al., 2001; Rekhter et al., 2019; Torrens-Spence et al., 2019), encode enzymes directly involved in SA biosynthesis. The type II genes, e.g., *PAD4* (Jirage et al., 1999), affect SA accumulation yet they do not have distinct enzymatic signatures. The biochemical function of the type II genes are largely unknown. The type III genes, e.g., *NPR1* (Wu et al., 2012; Ding et al., 2018), serve as signaling components to perceive and transduce the SA signal. The lack of a recognizable enzymatic feature in the predicted FLK protein suggests that FLK is either a type II or type III SA gene. To better understand how the *FLK* gene interacts genetically with some known SA pathway genes, we conducted genetic analyses, using the defense-sensitized mutant *acd6-1*. We crossed the *flk-1* mutant with several SA pathway mutants, including two type I SA mutants (*ics1-1* and *win3-1*), a type II SA mutant (*pad4-1*), and a type III mutant (*npr1-1*), in the *acd6-1* background. We made inferences on the interactions of two genes on the basis of the phenotypes of the triple mutant. A nonadditive suppression of *acd6-1-*conferred phenotypes would indicate that *FLK* and an SA gene act in the same pathway, whereas an additive suppression would suggest that the two genes function in separate pathways, or one is partially dependent on the other. This analysis has been successfully used to interrogate the genetic interactions among several defense genes (Song et al., 2004; Ng et al., 2011; Wang et al., 2011a; Wang et al., 2014; Hamdoun et al., 2016; Zhang et al., 2019). We found that all triple mutants containing *acd6-1flk-1* showed much larger plant stature than the corresponding double mutants (Figure 4A), suggesting that *flk-1* acts partially through or independently of these SA regulators in affecting *acd6-1-*conferred dwarfism. For cell death severity, all double mutants showed reduced cell death relative to *acd6-1* (Figure 4B and (Ng et al., 2011; Wang et al., 2011a)). We observed that all triple mutants had reduced cell death, compared with the corresponding double mutants, suggesting that *FLK* acts partially through or independently of the SA pathway genes, *ICS1, PAD4, WIN3*, and *NPR1*, in cell death regulation.

**Figure 4.**
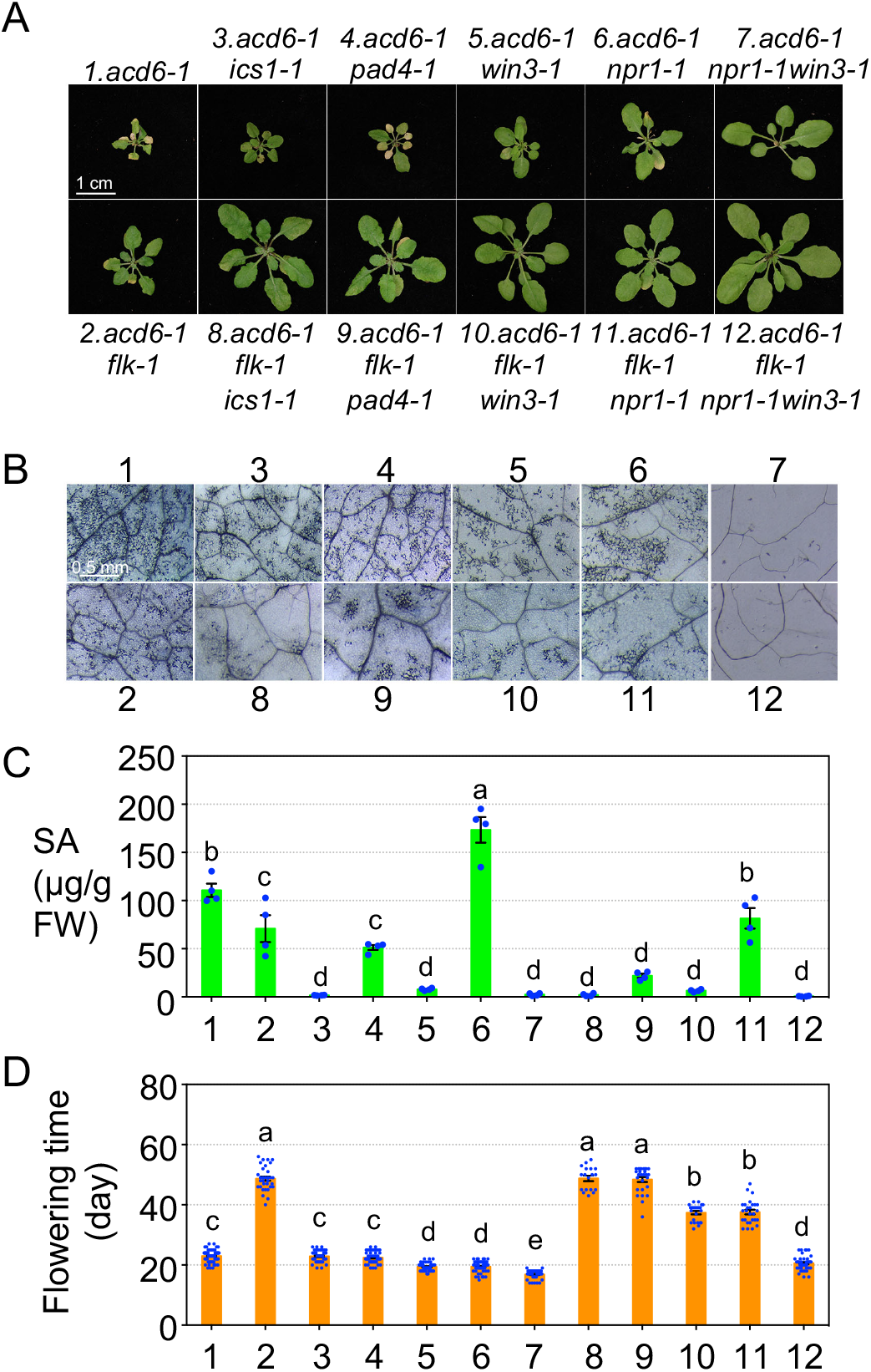
Genetic analysis of the interactions between *flk-1* and major SA mutants in *acd6-1*. (A) Pictures of 25-d old plants. The size bar represents 1 cm and is applied to all panels. (B) Cell death staining. The size bar represents 0.5 mm and is applied to all panels. (C) Total SA measurement by HPLC. (D) Flowering time. Error bars represent mean ± SEM in (C) (n=4) and (D) (n≥18). Different letters indicate significant difference among the samples (P<0.05; One-way ANOVA with post-hoc Tukey HSD test). Data for Col-0 and *flk-1* were shown in Figure 1. These experiments were repeated two times with similar results.

We further measured the total SA level in these plants (Figure 4C). Consistent with the roles of *ICS1* and *WIN3* as major SA synthesis genes, there was residual SA levels in *acd6-1ics1-1* and *acd6-1win3-1* with or without the presence of *flk-1*, suggesting that *FLK* acts fully through these SA synthesis genes in regulating SA accumulation. Further reduced SA levels were found in *acd6-1flk-1pad4-1* and *acd6-1flk-1npr1-1*, compared with the corresponding double mutants, suggesting that *FLK* acts partially through or independently of *PAD4* and *NPR1* in SA regulation. It is worth noting that NPR1 is an SA receptor that is known to both negatively and positively regulate SA levels, depending on the genetic composition of a plant (Ng et al., 2011; Wu et al., 2012). Consistent with this notion, *acd6-1npr1-1* accumulated a very high SA level, which was suppressed by *flk-1*. Because most triple mutants had a basal level of SA yet they did not show a complete suppression of plant size and cell death, additional SA-independent pathway(s) could contribute to these phenotypes.

### *FLK* regulation of flowering is independent of SA

For flowering time, we observed that *acd6-1, ics1-1*, and *pad4-1* did not affect delayed flowering in *flk-1* although these mutations drastically affected the SA level (Figure 4D). Thus, these results suggest that *FLK*-mediated flowering control is SA-independent. However, two other SA mutations, *npr1-1* and *win3-1*, suppressed late flowering in *flk-1*. We showed previously that *npr1-1* and *win3-1* work additively to stimulate early flowering and suppress *acd6-1-*conferred phenotypes (Wang et al., 2011a). Consistent with this observation, we found here that *npr1-1* and *win3-1* together completely suppressed *flk-1*-conferred late flowering, cell death, and reverted the plant stature to the wild type level in the quadruple mutant *acd6-1flk-1npr1-1win3-1*, compared with *acd6-1flk-1npr1-1* and *acd6-1flk-1win3-1* (Figure 4). The early flowering phenotype observed in *npr1-1* is likely due to a second site mutation in the background (Dong XN, personal communication). Additional loss-of-function alleles of *WIN3* were also associated with early flowering (Kenichi Tsuda and Yuelin Zhang, personal communications). However, whether this flowering regulatory function of *WIN3* is coupled with its role in SA biosynthesis remains to be determined. Together, these results suggest that *WIN3, NPR1*, and an unknown gene(s) act downstream of *FLK* to contribute to the regulation of flowering and *acd6-1-*conferred phenotypes, including SA accumulation, cell death, and plant size.

### RNA-seq analysis supports the role of *FLK* in defense regulation

To further elucidate how *FLK* is mechanistically linked to plant defense, we performed RNA-seq analysis. We generated transcriptome profiles of Col-0, *flk-1, flk-5, acd6-1, acd6-1flk-1*, and *acd6-1flk-5*, using the Illumina NovaSeq 6000 platform. Principal component analysis of the sequencing data after removing low-quality reads showed a correlation of gene expression profiles of the replicates in each genotype (Figure 5A). To identify *FLK*-affected genes, we compared four groups: (a) Col-0 vs. *flk-1*; (b) Col-0 vs. *flk-5*; (c) *acd6-1* vs. *acd6-1flk-1*; and (d) *acd6-1* vs. *acd6-1flk-5*. We found that there were a total of 8,083 genes differentially expressed (DE) in at least one pairwise comparison (adjusted p-value < 0.05) (Supplemental Data 1). Consistent with the known role of *FLK* in regulating development, Gene Ontology (GO) analyses showed that the DE genes were significantly enriched in those involved in primary metabolism (GO:0009415 and GO:0010243) and development-related processes (GO:0009639 and GO:0048511) (Figure 5B). The flowering regulator *FLC*, a known FLK target gene (Ripoll et al., 2009), is among the development-related DE genes affected by the *flk* mutations. In addition, a large number of defense genes and genes responding to abiotic stress were also found among the *FLK*-affected DE genes. A total of 955 defense genes showed differential expression in at least one comparison group (Figure 5C and Supplemental Data 1). Defense genes affected by *flk* mutants are from multiple signaling pathways involving SA, JA, ethylene (ET), programmed cell death (PCD), and/or PTI (Figures 5C and 5D). Expression of most of these defense genes was suppressed by the *flk* mutations in the *acd6-1* background, consistent with the suppression of *acd6-1-*conferred phenotypes by *flk*. For instance, in the SA pathway, SA biosynthesis genes (*ICS1*/*EDS16* and *PBS3/WIN3*), SA regulatory genes (*EDS1, PAD4, SAG101, CBP 60G*, and *SARD1*) (Feys et al., 2001; Feys et al., 2005; Wang et al., 2009; Zhang et al., 2010c; Wang et al., 2011b), and SA receptors (*NPR1* and *NPR3*) (Fu et al., 2012) showed lower expression in the *acd6-1flk* mutants than in *acd6-1* (Figure 5D and Supplemental Data 1). These data strongly support the role of *FLK* in SA regulation. In the JA pathway, we found that a number of genes involved in JA biosynthesis showed reduced expression in *acd6-1flk-1* and *acd6-1flk-5*, compared with *acd6-1*. These JA genes include the *LOX* genes (Vellosillo et al., 2007; Seltmann et al., 2010), *AOC3* (Hofmann and Pollmann, 2008), the *OPR* genes (Schaller et al., 2000; Stintzi and Browse, 2000), and several *JAZ* genes (Vanholme et al., 2007). However, we did not detect an expression change for the JA receptor gene *COI1*, the key downstream signal transducer *MYC2*, and several MYC2 target genes, including *ANAC019, ANAC055*, and *RD26* (Bu et al., 2008; Zheng et al., 2012). It is possible that only some branches of JA signaling are affected by the *flk* mutations. Additionally, we observed a downregulation of PTI-related genes in the *acd6-1flk* mutants, compared with *acd6-1*. These PTI genes include the pattern recognition receptors (*FLS2* and *EFR*), the co-receptor *BAK1*(Boller and Felix, 2009), PTI signal transducers (*MPK3, MKK5*, the *WRKY* genes (e.g., *WRKY22, WRKY* 29, *WRKY* 33, and other members in the family)) (Asai et al., 2002; Hsu et al., 2013), and members of the *ATL* gene family that show early responses to PTI (Libault et al., 2007; Lin et al., 2008). These data strongly support the role of *FLK* in PTI regulation.

**Figure 5.**
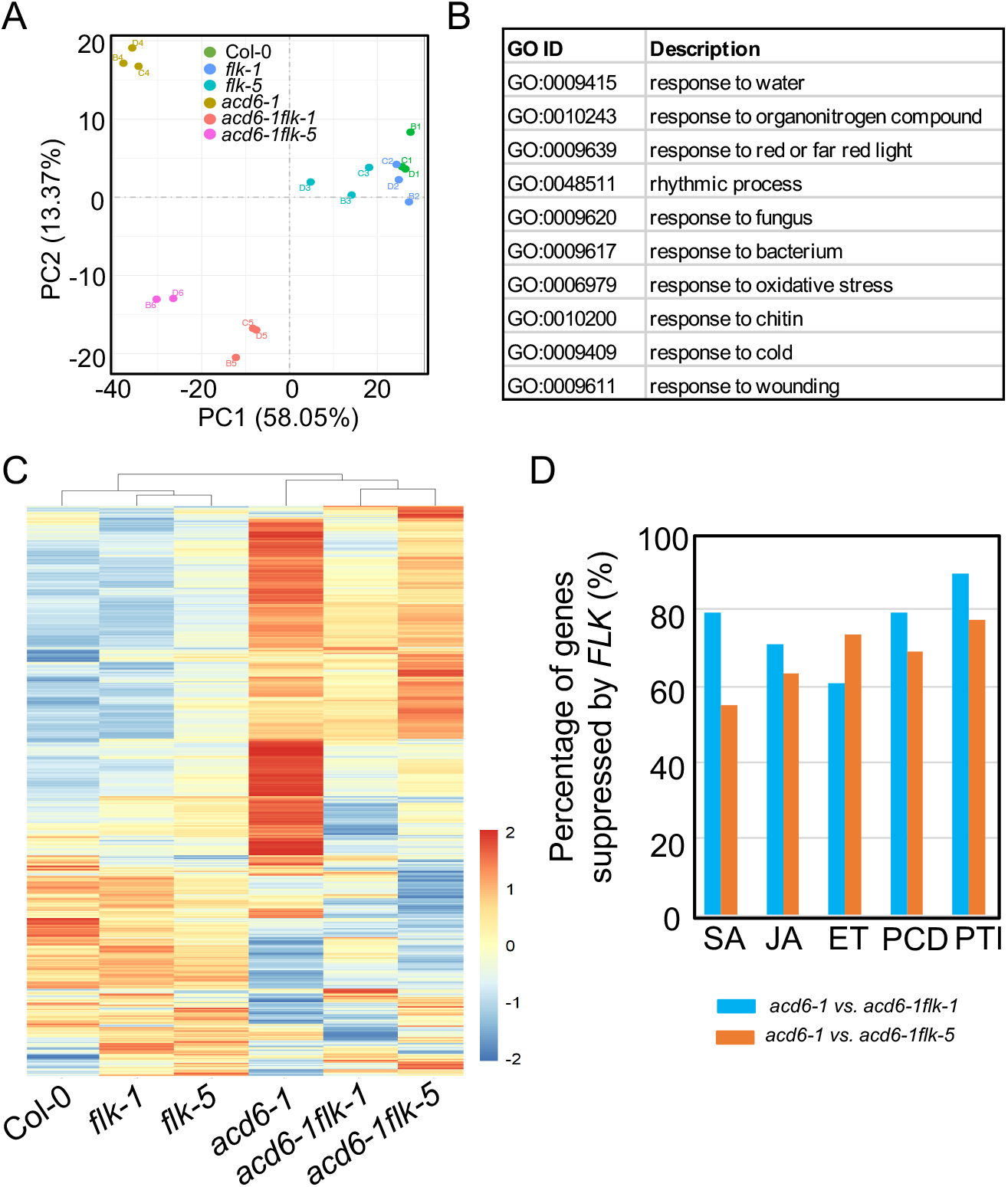
RNA-seq analysis reveals genes differentially affected by the *flk* mutations. (A) Principal component analysis. (B) Gene Ontology (GO) analysis of genes differentially affected by the *flk* mutations. GO categories were based on biological processes and they are significantly different in all four comparison groups: Col-0 vs. *flk-1*; (b) Col-0 vs. *flk-5*; (c) *acd6-1* vs. *acd6-1flk-1*; and (d) *acd6-1* vs. *acd6-1flk-5*. (C) Heatmap analysis of defense genes differentially affected by the *flk* mutations. (D) Suppression of major defense genes by the *flk* mutations in *acd6-1*. The GO category for SA is GO:0009751 (response to salicylic acid), for JA is GO:0009753 (response to jasmonic acid), for ethylene (ET) is GO:0009723 (response to ethylene), for programmed cell death (PCD) is GO:0008219 (cell death), and for PTI is GO:0007166 (cell surface receptor signaling pathway).

### *FLK* plays a role in ROS scavenging

While our data clearly support the roles of *FLK* in regulating SA and JA signaling, such function of *FLK* does not fully explain *flk-*conferred *Botrytis* resistance. This is because although the SA regulatory role of *FLK* supports *flk*-conferred *P. syringae* susceptibility, the decreased SA signaling did not necessarily lead to increased JA signaling in the *acd6-1lflk* mutants because like major SA genes, many JA biosynthesis genes are also down-regulated by the *flk* mutations (Figure 5D). In addition, the *flk* mutants showed a wild type-like response to MJ treatment (Figure 3C). In addition to SA and JA signaling, prior studies showed a correlation of reduced ROS with resistance to *Botrytis* and susceptibility to *P. syringae* in some studies (Chamnongpol et al., 1998; Govrin and Levine, 2000; Muckenschnabel et al., 2001; Polidoros et al., 2001). *flk-*conferred pathogen defense is consistent with compromised ROS accumulation and/or signaling. GO analyses of the RNA-seq data showed that genes regulated by the *flk-1* and *flk-5* mutations were significantly enriched in the oxidative stress category (Figure 5B). Thus, we investigated the link between *FLK* and ROS-mediated defense. The *flk* mutants had reduced ROS levels on the basis of these two experiments. First, the *flk* mutants showed reduced ROS burst upon elicitation of flg22 and elf26, compared with Col-0 and the two complementation lines (Figure 2D). Second, the high SA level in *acd6-1* was associated with an increase of H_2_O_2_, which was suppressed by the *flk* mutations (Supplemental Figure 2). Thus, *FLK* might play a role in regulating ROS homeostasis via ROS production and/or scavenging.

To further test how *FLK* regulates ROS, we treated plants with the herbicide paraquat, a redox-cycling agent that induces ROS production in a non-enzymatic manner in the chloroplast. Paraquat accepts electrons from photosystem I and transfers them to oxygen to produce ROS, subsequently interfering with photosynthetic electron transfer and leading to cell death (Lascano et al., 2012). Plant resistance to paraquat is often associated with increased ROS scavenging. Using methyl viologen (MV), a form of paraquat, we found that the *flk* mutants were less sensitive to MV treatment, displaying reduced leaf wilting, compared with Col-0 (Figures 6A and 6B). We further corroborated this result with the ion leakage assay, which reflects the severity of cell death induced by MV (Figure 6C). Together these data suggest a role of *FLK* in ROS scavenging.

**Figure 6.**
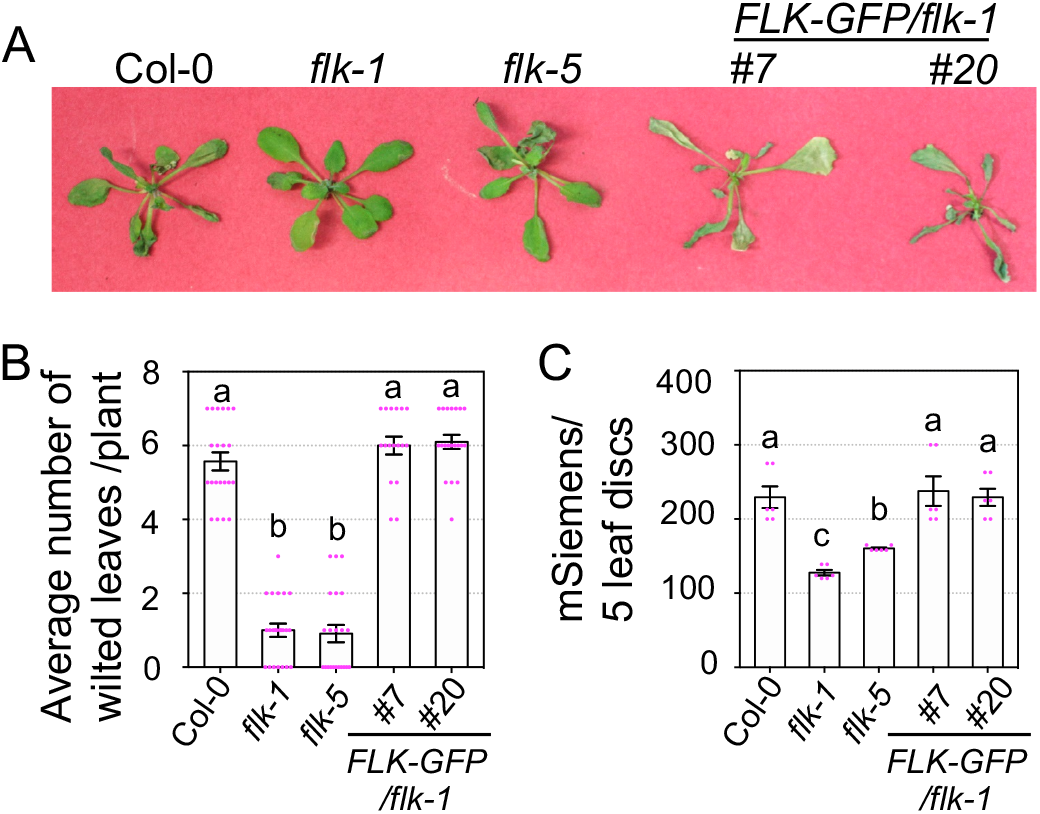
The *flk* mutations confer decreased MV sensitivity. (A) Images of plants treated with 40 µM MV. (B) The average number of wilted leaves in individual plants of each genotype 24 h after MV treatment. Mock treated plants were healthy and had no morphological difference (not shown). (C) Ion leakage assay. Discs from leaves treated with MV for 24 h were floated on water and measured for ion leakage. Error bars represent mean ± SEM in (B) (n=20) and (C) (n=3). Different letters indicate significant difference among the samples (P<0.05; One-way ANOVA with post-hoc Tukey HSD test). These experiments were repeated two times with similar results.

Some abiotic stress conditions, such as high salinity and UV, induce high ROS accumulation, causing oxidative damage and eventually cell death. Some plants resistant to abiotic stress are also more tolerant to MV (Kurepa et al., 1998; Tsugane et al., 1999; Overmyer et al., 2000; Fujibe et al., 2004), likely due to elevated ROS scavenging. We found that the *flk* mutants showed less chlorosis than Col-0 and *FLK-GFP* lines #7 and #20 under salinity, heat, or UV stress conditions (Supplemental Figure 3), suggesting a higher resistance of the *flk* mutants to these stress conditions. These results further support the role of *FLK* in ROS scavenging.

Many ROS scavenging enzymes detoxify ROS in the cell (Noctor et al., 2016; Smirnoff and Arnaud, 2019). To begin to address which enzymes are affected by *FLK*, we treated plant leaves with sodium azide (NaN_3_) to inhibit peroxidase activity followed by DAB staining for H_2_O_2_ accumulation (Lai et al., 2012). As expected, NaN_3_ treated Col-0 accumulated more H_2_O_2_ than mock treated plants (Figure 7A). If *FLK* acts through peroxidase in scavenging ROS, we expect that inhibition of peroxidase activity in the *flk* mutants would lead to higher ROS accumulation, compared with mock-treated plants. However, the *flk* mutants were less sensitive to NaN_3_ treatment and showed similar DAB staining as mock-treated samples. Thus, *FLK* unlikely acts through peroxidases in ROS scavenging. Similarly, we conducted a pharmacological inhibition assay, using 3-amino-1,2,4-triazole (3-AT) to inhibit the activity of catalases, another major class of ROS scavenging enzymes (Bestwick et al., 1997; Lai et al., 2012). We found that the *flk* mutants showed wild type-like DAB staining (Figure 7B), suggesting the importance of catalases in *FLK*-mediated ROS scavenging. qRT-PCR experiments further showed that expression of two catalase genes, *CAT2* and *CAT3*, was higher in the *flk* mutants (Figure 7B). Thus, *FLK* might act through catalases in ROS scavenging.

**Figure 7.**
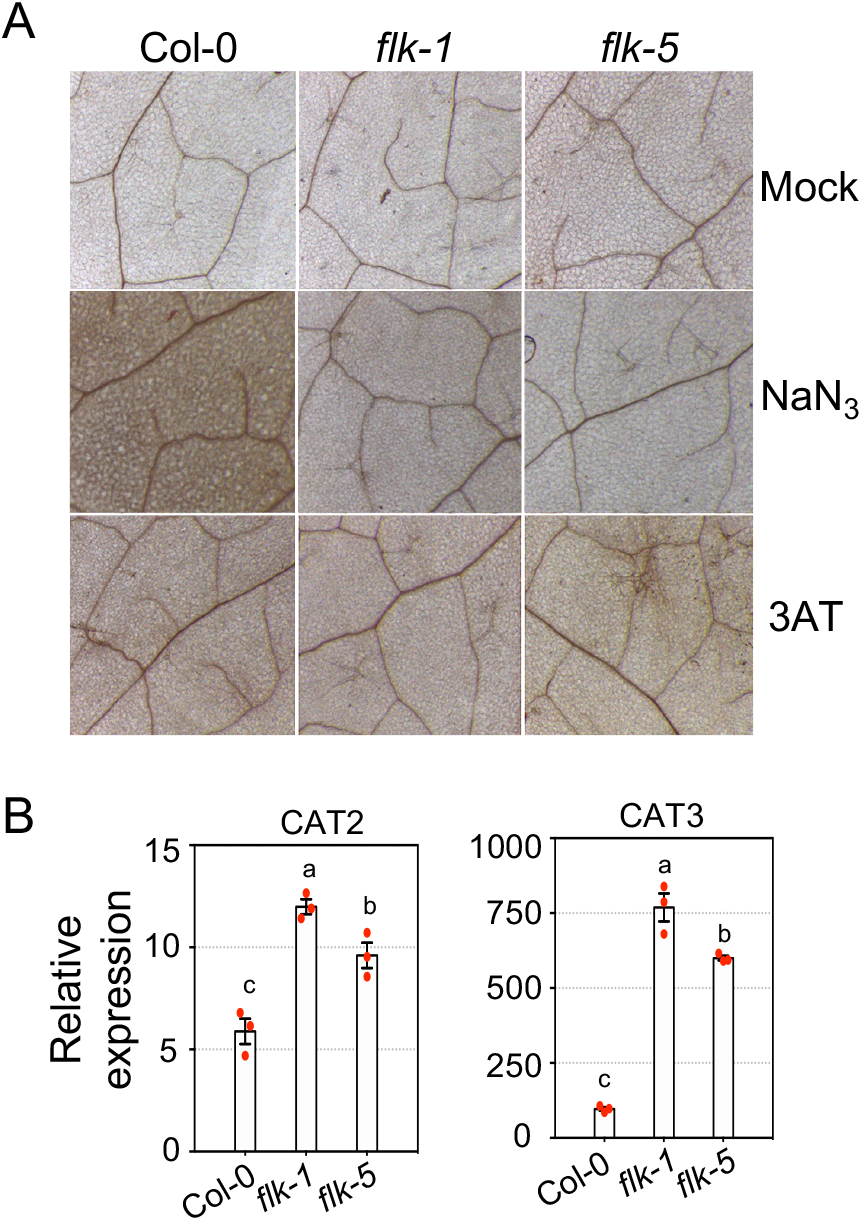
*FLK* likely acts through catalases in ROS scavenging. (A) DAB staining. Plants were treated with 2 mM NaN_3_, 10 mM 3-amino-1,2,4-triazole (3-AT), or mock solution for 24 h and were stained with DAB for H_2_O_2_ accumulation (brown staining).(B) qRT-PCR for gene expression. These experiments were repeated three times with similar results.

## Discussion

The *FLK* gene is known as a positive regulator of flowering time. We report here the identification of a new allele of *FLK* from a genetic screen aimed to uncover novel defense genes. The *flk* mutations conferred enhanced disease susceptibility to the biotroph *P. syringae* and enhanced disease resistance to the necrotroph *Botrytis*. Our data for the first time mechanistically link the *FLK* function with defense signaling and ROS scavenging. Thus, FLK is a multifunctional protein important for defense and development of plants.

In response to pathogens of varying lifestyles, plants activate downstream signaling pathways, some of which are antagonistic to one another. SA is a key signaling molecule critical for defense against biotrophs. SA might adversely affect plant defense against necrotrophs through crosstalking with other defense pathways, such as those mediated by JA and ROS. We showed here that the *flk* mutations suppressed high SA accumulation induced by *PmaDG3* infection and in the *acd6-1* background (Figures 1 and 2B). Genetic analysis revealed that *FLK*-regulated SA accumulation was fully dependent on the two major SA synthase genes *ICS1* and *WIN3* but partially dependent on *PAD4* and *NPR1* (Figure 4). RNA-seq analysis further showed that expression of most major SA regulatory genes were affected by the *flk* mutations. Together, these data strongly establish a role of *FLK* in SA regulation.

Because SA and JA are known to act antagonistically to defend against the biotrophic pathogen *P. syringae* and the necrotrophic pathogen *Botrytis*. The opposing response of the *flk* mutants to these two pathogens prompted us to investigate whether JA signaling is activated by *flk*. However, our data did not support an activation of the JA pathway because expression of many JA biosynthesis genes was suppressed by the *flk* mutations and the *flk* mutants behaved similarly to Col-0 in response to MJ treatment (Figures 3C and 5D). In addition to SA and JA signaling, prior studies showed a correlation of reduced ROS with resistance to *Botrytis* and susceptibility to *P. syringae* in some studies (Chamnongpol et al., 1998; Govrin and Levine, 2000; Muckenschnabel et al., 2001; Polidoros et al., 2001). Indeed, we found an association of the *flk* mutations with reduced ROS levels (Figure 2D and Supplemental Figure 2). Further experiments revealed that the *flk* mutants showed increased resistance to the ROS generating reagent MV, as well as stress imposed by UV, heat, or salinity, compared with Col-0 (Figure 6 and Supplemental Figure 2). These abiotic stress conditions are known to activate the oxidative response in plants and resistance to these conditions were shown to be related to enhanced ROS scavenging (Kurepa et al., 1998; Tsugane et al., 1999; Overmyer et al., 2000; Fujibe et al., 2004). Thus, these data together support the role of *FLK* in ROS scavenging.

ROS can be induced rapidly upon pathogen attack and abiotic stress conditions. ROS are also by-products of photosynthesis and respiration and are important for cellular communication and development in plants (Gapper and Dolan, 2006). While known as defense signaling molecules, high levels of ROS are toxic to plant cells. The multifunctional role of ROS makes it particularly important to balance ROS homeostasis in plant cells, which is not only controlled by numerous genes involved in ROS production but also by those involved ROS scavenging, including genes encoding catalases, ascorbate peroxidases, superoxide dismutases, peroxiredoxins, and other classes of peroxidases (Mittler et al., 2004; Smirnoff and Arnaud, 2019). Among the ROS scavenging enzymes, catalases are the most studied and are linked to pathogen defense (Mhamdi et al., 2010). Increased catalase activity, which results in decreased ROS, is associated with enhanced susceptibility to biotrophic pathogens but enhanced resistance to necrotrophic pathogens (Chamnongpol et al., 1998; Govrin and Levine, 2000; Polidoros et al., 2001; Rossi et al., 2017; Yuan et al., 2017). The *flk-*conferred responses to *P. syringae* (a biotroph) and *Botrytis* (a necrotroph) are consistent with reduced ROS levels and increased catalase gene expression in the *flk* mutants (Figures 2D, 7B, and Supplemental Figure 2). A pharmacological inhibition assay with a catalase inhibitor further supports the importance of catalases in *FLK*-mediated ROS scavenging (Figure 7A). Although peroxidase may not be required for *FLK* function in ROS scavenging, our data do not rule out that other types of ROS scavenging enzymes could also participate in *FLK*-mediated ROS homeostasis. In addition to a direct regulation of expression of catalase genes by *FLK*, it is possible that the increased ROS scavenging in *flk* is due to the crosstalk between SA and ROS signaling. High SA levels were shown to suppress ROS scavenging enzyme activities, including those of catalases and ascorbate peroxidases (Chen et al., 1993; Sanchez-Casas and Klessig, 1994; Durner and Klessig, 1995; Yuan et al., 2017). The suppression of SA by *flk* mutations could lead to a de-repression of catalase activity and subsequently higher ROS scavenging.

In addition to SA and ROS regulation, our data showed that the *flk* mutations compromised PTI upon elicitation with PAMP molecules and suppressed cell death in *acd6-1*. RNA-seq data revealed that major defense signaling genes are regulated by *FLK*, further providing mechanistic links of *FLK* function in pathogen defense. However, how the defense role of *FLK* is linked with its regulation of flowering time remains to be elucidated. Changes of SA levels have been shown to be associated with altered flowering time. For instance, exogenous application of SA led to early flowering in some plants (Endo et al., 2009; Wada et al., 2009). In Arabidopsis, a number of mutants showed an association of SA levels with early flowering (Martinez et al., 2004; Jin et al., 2008; March-Diaz et al., 2008; Liu et al., 2012). These observations underscore the importance of SA signaling in flowering control. In contrast to this notion, our data showed that *FLK-*mediated flowering is largely independent of SA level (Figure 4). Whether other defense signaling regulated by *FLK* is involved in flowering regulation remains to be elucidated. It is also possible that *FLK*’s role in flowering control might be through a different set of genes, independent of its role in defense regulation.

Although our data showed *FLK*-mediated flowering is largely independent of SA, we observed the suppression of late flowering in *flk* by two SA mutants, *npr1-1* and *win3-1*. Because the flowering time phenotype in the *npr1-1* mutant is likely due to a second site mutation, a further identification of this mutation will shed lights on the molecular pathway mediated by *FLK* in flowering control. The early flowering phenotype was also observed with other *WIN3* alleles (Kenichi Tsuda and Yuelin Zhang, personal communications), suggesting that *WIN3* plays roles in both flowering and SA biosynthesis. *WIN3* encodes a GH3 acyl adenylase-family enzyme that conjugates glutamate to isochorismate to produce isochorismate-9-glutamate, a precursor of SA (Rekhter et al., 2019; Torrens-Spence et al., 2019). It is possible that *WIN3* is not directly involved in flowering regulation. Instead, the lack of WIN3 enzymatic activity in the *win3-1* mutant might produce altered metabolic profile; one or more such metabolites could directly contribute to flowering regulation. Alternatively, *WIN3* has additional uncovered function independent of its role in SA synthesis for flowering regulation. *WIN3* suppresses expression of the flowering activator gene *FT* and promotes the repressor gene *FLC* (Wang et al., 2011a). This action of *WIN3* is opposite to that of *FLK* in regulating expression of these two genes (Lim et al., 2004; Mockler et al., 2004), forming the basis of *win3* suppression of *flk* in flowering control.

How could *FLK* execute its function in regulating various downstream pathways to affect defense and development of plants? *FLK* encodes a putative protein with three KH motifs. The KH motif is a highly conserved RNA binding motif found in proteins in diverse organisms. KH domain proteins were shown to affect RNA metabolism, such as pre-mRNA processing, mRNA stability, RNA transport, and/or translational efficiency (Nicastro et al., 2015). The KH domain proteins can function through a direct binding to, and/or an association in protein complexes with RNA molecules for processing (Nicastro et al., 2015). Consistent with *flk-*conferred development and defense phenotypes, our RNA-seq data revealed that a large number of genes involved in development and defense signaling were differentially expressed in the *flk* mutants, compared with control plants. These observations support the biochemical function of the FLK protein in regulating mRNA stability. It is possible that FLK directly binds to its target gene transcripts to influence their mRNA stability. Alternatively, FLK forms protein complexes with other proteins to regulate mRNA stability. In addition, FLK and/or its protein complexes could affect RNA alternative splicing. Indeed, two of the most similar human homologs of FLK, the PCBP and NOVA proteins, were shown to regulate RNA alternative splicing (Zhang et al., 2010a; Zhang et al., 2010b). Like these human homologs, FLK was reported to influence RNA alternative splicing of the target genes *FLC* and such a regulation resulted in the change of overall mRNA abundance (Ripoll et al., 2009). While the detailed biochemical mechanism by which FLK acts to regulate RNA metabolism remains to be elucidated, our data that show expression of many genes is affected by FLK and the lack of *FLK* results in altered defense and development, strongly support the multifunctionality of *FLK*. It is possible that such a multifunctionality of FLK can be decoupled by various downstream genes with distinct roles in defense and/or development regulation. Thus, it is critically important to uncover direct gene targets of FLK in order to further advance our understanding of the mechanistic actions of FLK.

Plant disease resistance is a complex process maintained in an intricate balance with normal development. Increasing evidence indicates the importance of post-transcriptional regulation of plant defense by RNA binding proteins (Lee and Kang, 2016). Our study supports the importance of FLK, a protein containing a conserved RNA binding motif, in plant development and defense. Because plants can activate conflicting defense strategies to fend off pathogens of different lifestyles, cautions should be taken when referring to the tradeoff between defense and development. Enhanced defense against a certain type of pathogens does not necessarily result in the cost to development, as demonstrated by the case to show *flk*-conferred opposing resistance to different pathogens and *flk*-conferred late flowering. It remains critically important to elucidate the detailed molecular mechanisms of gene function in regulating various physiological outputs. We expect that some FLK downstream targets decouple distinct FLK functions in defense and development. Such FLK pathway genes are potentially powerful molecular tools that can be used to develop novel biotechnological strategies to precisely control crop traits, maximizing plant growth while maintaining proper responses to environmental assaults.

## Materials and Methods

### Plant materials

Plants were grown in growth chambers with 180 µmol m^−2^ s^−1^ photon density, 60% humidity, and 22°C, and a photoperiod of 12 h light/12 h dark unless otherwise indicated. The *flk-1* mutant line (SALK_007750) was previously described (Mockler et al., 2004). *flk-5* was identified from the *acd6-1* suppressor screen (Lu et al., 2009). The *acd6-1flk-1* double mutant was generated by crossing the two single mutants and selecting the homozygous double mutant in the F2 generation, using proper primers. The triple mutants of *acd6-1flk-1* in combination with SA mutants, including *ics1-1, pad4-1, npr1-1*, and *win3-1*, were created by crossing the relevant mutants together. The triple mutants *acd6-1flk-1npr1-1* and *acd6-1flk-1win3-1* were previously described (Wang et al., 2011a) and they were used to generate the quadruple mutant *acd6-1flk-1npr1-1win3-1*. Primers for mutant allele detection were previously described (Ng et al., 2011; Wang et al., 2011a) or were listed in Supplemental Table 1.

### Generation of transgenic complementation lines

A fragment of 5.9 kb consisting of 2.4 kb *FLK* promoter and the 3.5 kb full*-*length *FLK* genomic fragment was PCR amplified and cloned in the pGlobug vector (Zhang and Mount, 2009), in frame at 3’ end with the reporter gene *GFP*. The *pFLK-FLK-GFP-NOS* fragment was subcloned into the binary vector pMLBart, using the NotI site. This construct was designated as *FLK-GFP* and was used to transform the *flk-1* mutant, using *Agrobacterium*-mediated plant transformation. Five homozygous transgenic plants that are independent transformants were obtained from the T2 generation.

### Pathogen infection

*Pseudomonas syringae pv. maculicola ES4326* strains DG3 (PmaDG3) was used to infect plants as previously described (Zhang et al., 2013). The fourth to seventh leaves of 25-d old plants were infiltrated with a PmaDG3 solution, using a needleless syringe. Discs of infected leaves were produced with a biopsy puncher of 4 mm diameter, homogenized in 10 mM MgSO4, and serially diluted. The dilutions were plated on King’s Broth agar media with 50 mg L^−1^ kanamycin for bacterial growth at 30 ºC.

*Botrytis cinerea* strain BO5-10 was kindly provided by Tesfaye Mengiste (Purdue University). A *Botrytis* solution of 2 × 10^5^ spores mL^−1^ was evenly sprayed onto plants for infection. Plants were covered for disease symptom development. Disease symptoms of the fourth to seventh leaves of each plant were scored three days post infection with the 1-5 scale as described (Wang et al., 2011a): 1 = no lesion or small rare lesions; 2 = lesions on 10% to 30% of a leaf; 3 = lesions on 30% to 50% of a leaf; 4 = lesions on 50% to 70% of a leaf; 5 = lesions on over 70% of a leaf.

### SA quantification

SA extraction and quantification were performed as previously described (Wang et al., 2011a). For pathogen-induced SA accumulation, the fourth to seventh leaves of 25-d old plants were infiltrated with 1 × 10^7^ cfu mL^−1^ PmaDG3 and were harvested at the indicated times for SA extraction. For non-infected plants, the whole plants were collected for SA extraction.

### Gene expression by qRT-PCR analysis

The fourth to seventh leaves of 25-d old plants were extracted for total RNA, using TRIzol reagent (Invitrogen) per manufacturer’s instructions. After the removal of genomic DNA, total RNA was reverse-transcribed, using a cDNA synthesis kit (ThermoFisher Scientific). Maxima SYBR Green/ROX qPCR Master Mix (ThermoFisher Scientific) was used for qPCR reactions. Primers for qRT-PCR were listed in Supplemental Table 1.

### Luminol assay

Leaf discs of 4 mm-diameter were isolated from the fourth to seventh leaves of 25-d old plants and floated atop 100 µL sterile water in a 96-well plate. The plate was covered with tin foil and placed in a growth chamber overnight for PAMP elicited ROS burst and for 2 h for basal ROS measurement. A 100 µL solution containing 300 µM L-012 (Wako Chemicals USA Inc. Richmond, VA) in 10 mM MOPS buffer (pH 7.4) and 1 µM flg22 or 1 µM elf26 was added to replace sterile water in each well and luminescence was recorded immediately using a Modulus II Microplate Reader.

### Cell death staining

Trypan blue staining was performed to visualize cell death as previously described (Ng et al., 2011). Briefly, the fourth to seventh leaves of 25-d old plants were harvested and boiled in lactophenol (phenol: glycerol: lactic acid: water=1:1:1:1; v/v) containing 0.01% trypan blue for 2 min. The stained leaves were boiled in alcoholic lactophenol (95% ethanol: lactophenol = 2:1) for 2 min, rinsed in 50% ethanol for 1 min, and finally cleared in 2.5g/ml chloral hydrate for 2 min. At least six leaves of each genotype were stained and visualized with a Leica M80 stereomicroscope. Cell death images were captured with a Leica IC80 HD camera connected to the microscope.

### Callose staining assay

The fourth to seventh leaves of 25-d old plants were infiltrated with 1 µM flg22 or sterile water. Leaves were harvested 24 h post infiltration and boiled in alcoholic lactophenol (2:1 95% ethanol: lactophenol) for 2 min followed by rinsing in 50% ethanol for 1 min. Cleared leaves were stained with aniline blue solution (0.01% aniline blue in 150 mM KH2PO4, pH 9.5) for 90 min in darkness. Callose deposition was visualized with a Leica M205FA fluorescence stereomicroscope and photographed with an Amscope CMOS digital camera.

### Root growth inhibition assays

Sterilized seeds were plated on agar plates containing 1/2 MS media supplemented with 1% sucrose (pH 5.7), stratified at 4º C for 2 days, and then transferred to a tissue culture chamber to grow for 4 days. Seedlings of relatively uniform size were transferred to a 24-well tissue culture plate with 1.5 mL 1 µM flg22, 10 µM MJ, or sterile water. A minimum of 4 seedlings per genotype per treatment were measured for root length 4 days post treatment and imaged with a Canon camera.

### MV sensitivity assay

Twenty-five-day old plants were sprayed with 40 µM methyl viologen (MV) or water as a control. At least 8 plants per genotype were recorded for leaf wilting 24 h post treatment. For ion leakage quantification, leaf discs from fourth to seventh leaves of plants were collected at 24 h post treatment and floated in 5 mL sterile water for 6 h followed by ion leakage recording, using a conductivity meter (Welwyn International Inc. Cleveland, OH, USA). Triplicate samples that each had five leaf discs were used for each genotype.

### Abiotic stress assays

Six-day old sterile seedlings grown on a plate with 1/2 MS media and 1% sucrose (pH 5.7) were used in the stress assays. For UV stress, seedlings were uncovered and placed under a Fisher Biotech transilluminator (312 nm) (Model FBTI-614; Fisher Scientific) for 2 h and then covered back and returned to a growth chamber. For heat stress, plates were sealed in plastic bags, submerged in a 44 ºC hot water bath for 4 hours, and then returned to a growth chamber. Images of seedlings treated with UV or heat stress were taken 48 h post treatment. For salinity stress, seedlings were placed in a 24-well plate containing 20 mM NaCl or water as a control and imaged seven days post treatment.

### Inhibition of ROS scavenging enzymes

Twenty-five-day old plants were sprayed with 2 mM NaN_3_, 10 mM 3-amino-1,2,4-triazole, or water as a control. The fourth to seventh leaves of the plants were collected 24 h post treatment followed by staining with a 3,3′-diaminobenzidine (DAB) solution as described (Daudi et al., 2012). Stained leaves were visualized with a Leica M205FA stereomicroscope and photographed with an Amscope CMOS digital camera.

### RNA-seq analysis

Total RNA was extracted from 25-d old Col-0, *flk-1, flk-5, acd6-1, acd6-1flk-1*, and *acd6-1flk-5* plants. A total amount of 1 μg RNA per sample was used to generate cDNA libraries, using NEBNext® UltraTM RNA Library Prep Kit for Illumina® (NEB, USA) following manufacturer’s recommendations. Deep sequencing was performed using Illumina NovaSeq 6000 (Novogene Corporation Inc.). Triplicate biological samples were used for all genotypes except *acd6-1flk-5*, which had duplicate samples because one sample failed to pass quality control. The samples were multiplexed and sequenced with the standard paired-end sequencing that has a read length of 150bp per end and 20M reads per end per sample. The raw reads in FASTQ format were filtered by removing reads containing adapters and reads of low quality and mapped to the Arabidopsis genome (TAIR 10). For global gene expression profiling, the relative expression value (Reads Per Kilobase of transcript per Million mapped reads (RPKM)) of higher than 0.3 was used as the cutoff to include genes for further analyses. Each RPKM value was corrected by adding the number one and then was log2-transformed. Differentially expressed genes in each comparison group were identified using the R package DESeq2, using the default parameters (Anders and Huber, 2010). Genes with an adjusted p-value < 0.05 found by DESeq2 were assigned as differentially expressed. Gene Ontology (GO) enrichment analysis of differentially expressed genes was implemented by the clusterProfiler R package, in which gene length bias was corrected. GO terms with corrected p-value less than 0.05 were considered significantly enriched by differential expressed genes. Graphical representation of gene expression correlation was analyzed, using heatmap.2 function.

## Acknowledgements

We thank the members in the Lu laboratory for their assistance in this work. This work was partially supported by a grant from National Science Foundation (NSF 1456140), UMBC Technology Catalyst Fund, and UMBC CENTRE Funding Initiative to HL.

## Author contributions

MF performed pathogen infections and PTI and ROS-related assays, measured flowering time and pathogen-induced SA accumulation, and assisted in the bioinformatics analysis. MG performed cell death staining, SA measurement, and gene expression analysis with the *flk* mutants, and prepared samples for the RNA-seq experiments. JS assisted in the cloning of the *FLK* gene. XZ cloned the *FLK-GFP* construct and conducted plant transformation and confocal microscopy for FLK-GFP localization. SK measured flowering time. PP assisted MF in PTI assays. AH assisted in the bioinformatics analysis. HL designed experiments, conducted genetic crosses, and wrote the manuscript with help from MF and MG.

## Additional Information

Competing financial interests: The authors declare no competing financial interests.

## Parsed Citations

Amasino, R. (2010). Seasonal and developmental timing of flowering. Plant J 61, 1001–1013.

Anders, S., and Huber, W. (2010). Differential expression analysis for sequence count data. Genome Biol 11, R106.

Asai, T., Tena, G., Plotnikova, J., Willmann, M.R., Chiu, W.L., Gomez-Gomez, L., Boller, T., Ausubel, F.M., and Sheen, J. (2002). MAP kinase signalling cascade in Arabidopsis innate immunity. Nature 415, 977–983.

Bari, R., and Jones, J.D. (2009). Role of plant hormones in plant defence responses. Plant Mol Biol 69, 473–488.

Bestwick, C.S., Brown, I.R., Bennett, M.H., and Mansfield, J.W. (1997). Localization of hydrogen peroxide accumulation during the hypersensitive reaction of lettuce cells to Pseudomonas syringae pv phaseolicola. Plant Cell 9, 209–221.

Boller, T., and Felix, G. (2009). Arenaissance of elicitors: perception of microbe-associated molecular patterns and danger signals by pattern-recognition receptors. Annu Rev Plant Biol 60, 379–406.

Bu, Q., Jiang, H., Li, C.B., Zhai, Q., Zhang, J., Wu, X., Sun, J., Xie, Q., and Li, C. (2008). Role of the Arabidopsis thaliana NAC transcription factors ANAC019 and ANAC055 in regulating jasmonic acid-signaled defense responses. Cell Res 18, 756–767.

Campos, M.L., Kang, J.H., and Howe, G.A. (2014). Jasmonate-triggered plant immunity. J Chem Ecol 40, 657–675.

Chamnongpol, S., Willekens, H., Moeder, W., Langebartels, C., Sandermann, H., Jr., Van Montagu, M., Inze, D., and Van Camp, W. (1998). Defense activation and enhanced pathogen tolerance induced by H_2_O_2_ in transgenic tobacco. Proc. Natl. Acad. Sci. USA 95, 5818–5823.

Chen, T., Cui, P., Chen, H., Ali, S., Zhang, S., and Xiong, L. (2013). AKH-domain RNA-binding protein interacts with FIERY2/CTD phosphatase-like 1 and splicing factors and is important for pre-mRNA splicing in Arabidopsis. PLoS Genet 9, e1003875.

Chen, Z., Silva, H., and Klessig, D.F. (1993). Active oxygen species in the induction of plant systemic acquired resistance by salicylic acid. Science 262, 1883–1886.

Cheng, Y., Kato, N., Wang, W., Li, J., and Chen, X. (2003). Two RNAbinding proteins, HEN4 and HUA1, act in the processing of AGAMOUS pre-mRNA in Arabidopsis thaliana. Dev Cell 4, 53–66.

Daudi, A., Cheng, Z., O’Brien, J.A., Mammarella, N., Khan, S., Ausubel, F.M., and Bolwell, G.P. (2012). The apoplastic oxidative burst peroxidase in Arabidopsis is a major component of pattern-triggered immunity. Plant Cell 24, 275–287.

Ding, Y., Sun, T., Ao, K., Peng, Y., Zhang, Y., Li, X., and Zhang, Y. (2018). Opposite Roles of Salicylic Acid Receptors NPR1 and NPR3/NPR4 in Transcriptional Regulation of Plant Immunity. Cell 173, 1454–1467 e1415.

Dodds, P.N., and Rathjen, J.P. (2010). Plant immunity: towards an integrated view of plant-pathogen interactions. Nat Rev Genet 11, 539–548.

Durner, J., and Klessig, D.F. (1995). Inhibition of ascorbate peroxidase by salicylic acid and 2,6-dichloroisonicotinic acid, two inducers of plant defense responses. Proc Natl Acad Sci U S A 92, 11312–11316.

Endo, J., Takahashi, W., Ikegami, T., Beppu, T., and Tanaka, O. (2009). Induction of flowering by inducers of systemic acquired resistance in the Lemna plant. Biosci Biotechnol Biochem 73, 183–185.

Feys, B.J., Moisan, L.J., Newman, M.A., and Parker, J.E. (2001). Direct interaction between the Arabidopsis disease resistance signaling proteins, EDS1 and PAD4. EMBO J. 20, 5400–5411.

Feys, B.J., Wiermer, M., Bhat, R.A., Moisan, L.J., Medina-Escobar, N., Neu, C., Cabral, A., and Parker, J.E. (2005). Arabidopsis SENESCENCE-ASSOCIATED GENE101 stabilizes and signals within an ENHANCED DISEASE SUSCEPTIBILITY1 complex in plant innate immunity. Plant Cell 17, 2601–2613.

Fu, Z.Q., and Dong, X. (2013). Systemic acquired resistance: turning local infection into global defense. Annu. Rev. Plant Biol. 64, 839–863.

Fu, Z.Q., Yan, S., Saleh, A., Wang, W., Ruble, J., Oka, N., Mohan, R., Spoel, S.H., Tada, Y., Zheng, N., and Dong, X. (2012). NPR3 and NPR4 are receptors for the immune signal salicylic acid in plants. Nature 486, 228–232.

Fujibe, T., Saji, H., Arakawa, K., Yabe, N., Takeuchi, Y., and Yamamoto, K.T. (2004). A methyl viologen-resistant mutant of Arabidopsis, which is allelic to ozone-sensitive rcd1, is tolerant to supplemental ultraviolet-B irradiation. Plant Physiol 134, 275–285.

Gapper, C., and Dolan, L. (2006). Control of plant development by reactive oxygen species. Plant Physiol 141, 341–345.

Genoud, T., Buchala, A.J., Chua, N.H., and Metraux, J.P. (2002). Phytochrome signalling modulates the SA-perceptive pathway in Arabidopsis. Plant J. 31, 87–95.

Geuens, T., Bouhy, D., and Timmerman, V. (2016). The hnRNP family: insights into their role in health and disease. Hum Genet 135, 851–867.

Glazebrook, J. (2005). Contrasting mechanisms of defense against biotrophic and necrotrophic pathogens. Annu Rev Phytopathol 43, 205–227.

Govrin, E., and Levine, A. (2000). The hypersensitive response facilitates plant infection by the necrotrophic pathogen Botrytis cinerea. Current Biol. 10, 751–757.

Griebel, T., and Zeier, J. (2008). Light regulation and daytime dependency of inducible plant defenses in Arabidopsis: phytochrome signaling controls systemic acquired resistance rather than local defense. Plant Physiol 147, 790–801.

Guan, Q., Wen, C., Zeng, H., and Zhu, J. (2013). AKH domain-containing putative RNA-binding protein is critical for heat stress-responsive gene regulation and thermotolerance in Arabidopsis. Mol Plant 6, 386–395.

Hamdoun, S., Zhang, C., Gill, M., Kumar, N., Churchman, M., Larkin, J.C., Kwon, A., and Lu, H. (2016). Differential roles of two homologous cyclin-dependent kinase inhibitor genes in regulating cell cycle and innate immunity in Arabidopsis. Plant Physiol 170, 515–527.

Herrera-Vásquez, A., Salinas, P., and Holuigue, L. (2015). Salicylic acid and reactive oxygen species interplay in the transcriptional control of defense genes expression. Frontiers in Plant Science 6.

Hofmann, E., and Pollmann, S. (2008). Molecular mechanism of enzymatic allene oxide cyclization in plants. Plant Physiol Biochem 46, 302–308.

Hsu, F.C., Chou, M.Y., Chou, S.J., Li, Y.R., Peng, H.P., and Shih, M.C. (2013). Submergence confers immunity mediated by the WRKY22 transcription factor in Arabidopsis. Plant Cell 25, 2699–2713.

Jin, J.B., Jin, Y.H., Lee, J., Miura, K., Yoo, C.Y., Kim, W.Y., Van Oosten, M., Hyun, Y., Somers, D.E., Lee, I., Yun, D.J., Bressan, R.A., and Hasegawa, P.M. (2008). The SUMO E3 ligase, AtSIZ1, regulates flowering by controlling a salicylic acid-mediated floral promotion pathway and through affects on FLC chromatin structure. Plant J 53, 530–540.

Jirage, D., Tootle, T.L., Reuber, T.L., Frost, L.N., Feys, B.J., Parker, J.E., Ausubel, F.M., and Glazebrook, J. (1999). Arabidopsis thaliana PAD4 encodes a lipase-like gene that is important for salicylic acid signaling. Proc. Natl. Acad. Sci. U. S. A. 96, 13583–13588.

Karasov, T.L., Chae, E., Herman, J.J., and Bergelson, J. (2017). Mechanisms to Mitigate the Trade-Off between Growth and Defense. Plant Cell 29, 666–680.

Korves, T.M., and Bergelson, J. (2003). Adevelopmental response to pathogen infection in Arabidopsis. Plant Physiol 133, 339–347.

Kunze, G., Zipfel, C., Robatzek, S., Niehaus, K., Boller, T., and Felix, G. (2004). The N terminus of bacterial elongation factor Tu elicits innate immunity in Arabidopsis plants. Plant Cell 16, 3496–3507.

Kurepa, J., Smalle, J., Van Montagu, M., and Inze, D. (1998). Oxidative stress tolerance and longevity in Arabidopsis: the late-flowering mutant gigantea is tolerant to paraquat. Plant J 14, 759–764.

Lai, A.G., Doherty, C.J., Mueller-Roeber, B., Kay, S.A., Schippers, J.H., and Dijkwel, P.P. (2012). CIRCADIAN CLOCK-ASSOCIATED 1 regulates ROS homeostasis and oxidative stress responses. Proc Natl Acad Sci U S A 109, 17129–17134.

Lascano, R., Munoz, N., Robert, G., Rodriguez, M., Melchiorre, M., Trippi, V., and Quero, G. (2012). Paraquat: An Oxidative Stress Inducer. InTech.

Lee, K., and Kang, H. (2016). Emerging Roles of RNA-Binding Proteins in Plant Growth, Development, and Stress Responses. Mol Cells 39, 179–185.

Li, L., Li, M., Yu, L., Zhou, Z., Liang, X., Liu, Z., Cai, G., Gao, L., Zhang, X., Wang, Y., Chen, S., and Zhou, J.M. (2014). The FLS2-associated kinase BIK1 directly phosphorylates the NADPH oxidase RbohD to control plant immunity. Cell Host Microbe 15, 329–338.

Libault, M., Wan, J., Czechowski, T., Udvardi, M., and Stacey, G. (2007). Identification of 118 Arabidopsis transcription factor and 30 ubiquitin-ligase genes responding to chitin, a plant-defense elicitor. Mol Plant Microbe Interact 20, 900–911.

Lim, M.H., Kim, J., Kim, Y.S., Chung, K.S., Seo, Y.H., Lee, I., Hong, C.B., Kim, H.J., and Park, C.M. (2004). Anew Arabidopsis gene, FLK, encodes an RNAbinding protein with K homology motifs and regulates flowering time via FLOWERING LOCUS C. Plant Cell 16, 731–740.

Lin, S.S., Martin, R., Mongrand, S., Vandenabeele, S., Chen, K.C., Jang, I.C., and Chua, N.H. (2008). RING1 E3 ligase localizes to plasma membrane lipid rafts to trigger FB1-induced programmed cell death in Arabidopsis. Plant J 56, 550–561.

Liu, J., Li, W., Ning, Y., Shirsekar, G., Cai, Y., Wang, X., Dai, L., Wang, Z., Liu, W., and Wang, G.L. (2012). The U-Box E3 ligase SPL11/PUB13 is a convergence point of defense and flowering signaling in plants. Plant Physiol 160, 28–37.

Lorkovic, Z.J., and Barta, A. (2002). Genome analysis: RNArecognition motif (RRM) and K homology (KH) domain RNA-binding proteins from the flowering plant Arabidopsis thaliana. Nucleic Acids Res 30, 623–635.

Lu, H. (2009). Dissection of salicylic acid-mediated defense signaling networks. Plant Signal. Behav. 4, 713–717.

Lu, H., McClung, C.R., and Zhang, C. (2017). Tick tock: circadian regulation of plant innate immunity. Annu Rev Phytopathol 55, 287–311.

Lu, H., Rate, D.N., Song, J.T., and Greenberg, J.T. (2003). ACD6, a novel ankyrin protein, is a regulator and an effector of salicylic acid signaling in the Arabidopsis defense response. Plant Cell 15, 2408–2420.

Lu, H., Salimian, S., Gamelin, E., Wang, G., Fedorowski, J., LaCourse, W., and Greenberg, J.T. (2009). Genetic analysis of acd6-1 reveals complex defense networks and leads to identification of novel defense genes in Arabidopsis. Plant J 58, 401–412.

Lyons, R., Iwase, A., Gansewig, T., Sherstnev, A., Duc, C., Barton, G.J., Hanada, K., Higuchi-Takeuchi, M., Matsui, M., Sugimoto, K., Kazan, K., Simpson, G.G., and Shirasu, K. (2013). The RNA-binding protein FPAregulates flg22-triggered defense responses and transcription factor activity by alternative polyadenylation. Sci Rep 3, 2866.

Macho, A.P., Boutrot, F., Rathjen, J.P., and Zipfel, C. (2012). Aspartate oxidase plays an important role in Arabidopsis stomatal immunity. Plant Physiol 159, 1845–1856.

March-Diaz, R., Garcia-Dominguez, M., Lozano-Juste, J., Leon, J., Florencio, F.J., and Reyes, J.C. (2008). Histone H2A.Z and homologues of components of the SWR1 complex are required to control immunity in Arabidopsis. Plant J 53, 475–487.

Martinez, C., Pons, E., Prats, G., and Leon, J. (2004). Salicylic acid regulates flowering time and links defence responses and reproductive development. Plant J 37, 209–217.

Mhamdi, A., Queval, G., Chaouch, S., Vanderauwera, S., Van Breusegem, F., and Noctor, G. (2010). Catalase function in plants: a focus on Arabidopsis mutants as stress-mimic models. J Exp Bot 61, 4197–4220.

Mittler, R., Vanderauwera, S., Gollery, M., and Van Breusegem, F. (2004). Reactive oxygen gene network of plants. Trends Plant Sci 9, 490–498.

Mockler, T.C., Yu, X., Shalitin, D., Parikh, D., Michael, T.P., Liou, J., Huang, J., Smith, Z., Alonso, J.M., Ecker, J.R., Chory, J., and Lin, C. (2004). Regulation of flowering time in Arabidopsis by K homology domain proteins. Proc Natl Acad Sci U S A 101, 12759–12764.

Muckenschnabel, I., Goodman, B.A., Deighton, N., Lyon, G.D., and Williamson, B. (2001). Botrytis cinerea induces the formation of free radicals in fruits of Capsicum annuum at positions remote from the site of infection. Protoplasma 218, 112–116.

Ng, G., Seabolt, S., Zhang, C., Salimian, S., Watkins, T.A., and Lu, H. (2011). Genetic dissection of salicylic acid-mediated defense signaling networks in Arabidopsis. Genetics 189, 851–859.

Nicastro, G., Taylor, I.A., and Ramos, A. (2015). KH-RNAinteractions: back in the groove. Curr Opin Struct Biol 30, 63–70.

Ning, Y., Liu, W., and Wang, G.L. (2017). Balancing Immunity and Yield in Crop Plants. Trends Plant Sci 22, 1069–1079.

Noctor, G., Mhamdi, A., and Foyer, C.H. (2016). Oxidative stress and antioxidative systems: recipes for successful data collection and interpretation. Plant Cell Environ 39, 1140–1160.

Ortuno-Miquel, S., Rodriguez-Cazorla, E., Zavala-Gonzalez, E.A., Martinez-Laborda, A., and Vera, A. (2019). Arabidopsis HUA ENHANCER 4 delays flowering by upregulating the MADS-box repressor genes FLC and MAF4. Sci Rep 9, 1478.

Overmyer, K., Tuominen, H., Kettunen, R., Betz, C., Langebartels, C., Sandermann, H., Jr., and Kangasjarvi, J. (2000). Ozone-sensitive arabidopsis rcd1 mutant reveals opposite roles for ethylene and jasmonate signaling pathways in regulating superoxide-dependent cell death. Plant Cell 12, 1849–1862.

Polidoros, A.N., Mylona, P.V., and Scandalios, J.G. (2001). Transgenic tobacco plants expressing the maize Cat2 gene have altered catalase levels that affect plant-pathogen interactions and resistance to oxidative stress. Transgenic Res 10, 555–569.

Rate, D.N., Cuenca, J.V., Bowman, G.R., Guttman, D.S., and Greenberg, J.T. (1999). The gain-of-function Arabidopsis acd6 mutant reveals novel regulation and function of the salicylic acid signaling pathway in controlling cell death, defenses, and cell growth. Plant Cell 11, 1695–1708.

Rekhter, D., Ludke, D., Ding, Y., Feussner, K., Zienkiewicz, K., Lipka, V., Wiermer, M., Zhang, Y., and Feussner, I. (2019). Isochorismate-derived biosynthesis of the plant stress hormone salicylic acid. Science 365, 498–502.

Ripoll, J.J., Rodriguez-Cazorla, E., Gonzalez-Reig, S., Andujar, A., Alonso-Cantabrana, H., Perez-Amador, M.A., Carbonell, J., Martinez-Laborda, A., and Vera, A. (2009). Antagonistic interactions between Arabidopsis K-homology domain genes uncover PEPPER as a positive regulator of the central floral repressor FLOWERING LOCUS C. Dev Biol 333, 251–262.

Roden, L.C., and Ingle, R.A. (2009). Lights, rhythms, infection: the role of light and the circadian clock in determining the outcome of plant-pathogen interactions. Plant Cell 21, 2546–2552.

Rossi, F.R., Krapp, A.R., Bisaro, F., Maiale, S.J., Pieckenstain, F.L., and Carrillo, N. (2017). Reactive oxygen species generated in chloroplasts contribute to tobacco leaf infection by the necrotrophic fungus Botrytis cinerea. Plant J 92, 761–773.

Sanchez-Casas, P., and Klessig, D.F. (1994). Asalicylic Acid-Binding Activity and a Salicylic Acid-Inhibitable Catalase Activity Are Present in a Variety of Plant Species. Plant Physiol 106, 1675–1679.

Schaller, F., Biesgen, C., Mussig, C., Altmann, T., and Weiler, E.W. (2000). 12-Oxophytodienoate reductase 3 (OPR3) is the isoenzyme involved in jasmonate biosynthesis. Planta 210, 979–984.

Seltmann, M.A., Stingl, N.E., Lautenschlaeger, J.K., Krischke, M., Mueller, M.J., and Berger, S. (2010). Differential impact of lipoxygenase 2 and jasmonates on natural and stress-induced senescence in Arabidopsis. Plant Physiol 152, 1940–1950.

Singh, V., Roy, S., Giri, M.K., Chaturvedi, R., Chowdhury, Z., Shah, J., and Nandi, A.K. (2013). Arabidopsis thaliana FLOWERING LOCUS D is required for systemic acquired resistance. Mol Plant Microbe Interact 26, 1079–1088.

Smirnoff, N., and Arnaud, D. (2019). Hydrogen peroxide metabolism and functions in plants. New Phytol 221, 1197–1214.

Song, J.T., Lu, H., McDowell, J.M., and Greenberg, J.T. (2004). Akey role for ALD1 in activation of local and systemic defenses in Arabidopsis. Plant J. 40, 200–212.

Spoel, S.H., and Dong, X. (2008). Making sense of hormone crosstalk during plant immune responses. Cell Host Microbe 3, 348–351.

Stintzi, A., and Browse, J. (2000). The Arabidopsis male-sterile mutant, opr3, lacks the 12-oxophytodienoic acid reductase required for jasmonate synthesis. Proc Natl Acad Sci U S A 97, 10625–10630.

Survila, M., Davidsson, P.R., Pennanen, V., Kariola, T., Broberg, M., Sipari, N., Heino, P., and Palva, E.T. (2016). Peroxidase-Generated Apoplastic ROS Impair Cuticle Integrity and Contribute to DAMP-Elicited Defenses. Front Plant Sci 7, 1945.

Thatcher, L.F., Kamphuis, L.G., Hane, J.K., Onate-Sanchez, L., and Singh, K.B. (2015). The Arabidopsis KH-Domain RNA-Binding Protein ESR1 Functions in Components of Jasmonate Signalling, Unlinking Growth Restraint and Resistance to Stress. PLoS One 10, e0126978.

Torrens-Spence, M.P., Bobokalonova, A., Carballo, V., Glinkerman, C.M., Pluskal, T., Shen, A., and Weng, J.K. (2019). PBS3 and EPS1 Complete Salicylic Acid Biosynthesis from Isochorismate in Arabidopsis. Mol Plant 12, 1577–1586.

Toruno, T.Y., Stergiopoulos, I., and Coaker, G. (2016). Plant-Pathogen Effectors: Cellular Probes Interfering with Plant Defenses in Spatial and Temporal Manners. Annu Rev Phytopathol 54, 419–441.

Tsugane, K., Kobayashi, K., Niwa, Y., Ohba, Y., Wada, K., and Kobayashi, H. (1999). Arecessive Arabidopsis mutant that grows photoautotrophically under salt stress shows enhanced active oxygen detoxification. Plant Cell 11, 1195–1206.

Vanholme, B., Grunewald, W., Bateman, A., Kohchi, T., and Gheysen, G. (2007). The tify family previously known as ZIM. Trends Plant Sci 12, 239–244.

Vellosillo, T., Martinez, M., Lopez, M.A., Vicente, J., Cascon, T., Dolan, L., Hamberg, M., and Castresana, C. (2007). Oxylipins produced by the 9-lipoxygenase pathway in Arabidopsis regulate lateral root development and defense responses through a specific signaling cascade. Plant Cell 19, 831–846.

Wada, K.C., Yamada, M., Shiraya, T., and Takeno, K. (2009). Salicylic acid and the flowering gene FLOWERING LOCUS T homolog are involved in poor-nutrition stress-induced flowering of Pharbitis nil. J Plant Physiol 167, 447–452.

Wang, G., Zhang, C., Battle, S., and Lu, H. (2014). The phosphate transporter PHT4;1 is a Salicylic acid regulator likely controlled by the circadian clock protein CCA1. Frontiers in Plant Science 5.

Wang, G.F., Seabolt, S., Hamdoun, S., Ng, G., Park, J., and Lu, H. (2011a). Multiple roles of WIN3 in regulating disease resistance, cell death, and flowering time in Arabidopsis. Plant Physiol. 156, 1508–1519.

Wang, L., Tsuda, K., Sato, M., Cohen, J.D., Katagiri, F., and Glazebrook, J. (2009). Arabidopsis CaM binding protein CBP60g contributes to MAMP-induced SAaccumulation and is involved in disease resistance against Pseudomonas syringae. PLoS Pathog 5, e1000301.

Wang, L., Tsuda, K., Truman, W., Sato, M., Nguyen le, V., Katagiri, F., and Glazebrook, J. (2011b). CBP60g and SARD1 play partially redundant critical roles in salicylic acid signaling. Plant J 67, 1029–1041.

Wildermuth, M.C., Dewdney, J., Wu, G., and Ausubel, F.M. (2001). Isochorismate synthase is required to synthesize salicylic acid for plant defence. Nature 414, 562–565.

Woloszynska, M., Le Gall, S., Neyt, P., Boccardi, T.M., Grasser, M., Langst, G., Aesaert, S., Coussens, G., Dhondt, S., Van De Slijke, E., Bruno, L., Fung-Uceda, J., Mas, P., Van Montagu, M., Inze, D., Himanen, K., De Jaeger, G., Grasser, K.D., and Van Lijsebettens, M. (2019). Histone 2B monoubiquitination complex integrates transcript elongation with RNAprocessing at circadian clock and flowering regulators. Proc Natl Acad Sci U S A 116, 8060–8069.

Wu, Y., Zhang, D., Chu, J.Y., Boyle, P., Wang, Y., Brindle, I.D., De Luca, V., and Despres, C. (2012). The Arabidopsis NPR1 protein is a receptor for the plant defense hormone salicylic acid. Cell Rep. 1, 639–647.

Yan, Z., Jia, J., Yan, X., Shi, H., and Han, Y. (2017). Arabidopsis KHZ1 and KHZ2, two novel non-tandem CCCH zinc-finger and Khomolog domain proteins, have redundant roles in the regulation of flowering and senescence. Plant Mol Biol 95, 549–565.

Yuan, H.M., Liu, W.C., and Lu, Y.T. (2017). CATALASE2 Coordinates SA-Mediated Repression of Both Auxin Accumulation and JA Biosynthesis in Plant Defenses. Cell Host Microbe 21, 143–155.

Zhang, C., Frias, M.A., Mele, A., Ruggiu, M., Eom, T., Marney, C.B., Wang, H., Licatalosi, D.D., Fak, J.J., and Darnell, R.B. (2010a). Integrative modeling defines the Nova splicing-regulatory network and its combinatorial controls. Science 329, 439–443.

Zhang, C., Xie, Q., Anderson, R.G., Ng, G., Seitz, N.C., Peterson, T., McClung, C.R., McDowell, J.M., Kong, D., Kwak, J.M., and Lu, H. (2013). Crosstalk between the circadian clock and innate immunity in Arabidopsis. PLoS Pathog. 9, e1003370.

Zhang, C., Gao, M., Seitz, N.C., Angel, W., Hallworth, A., Wiratan, L., Darwish, O., Alkharouf, N., Dawit, T., Lin, D., Egoshi, R., Wang, X., McClung, C.R., and Lu, H. (2019). LUX ARRHYTHMO mediates crosstalk between the circadian clock and defense in Arabidopsis. Nat Commun 10, 2543.

Zhang, T., Huang, X.H., Dong, L., Hu, D., Ge, C., Zhan, Y.Q., Xu, W.X., Yu, M., Li, W., Wang, X., Tang, L., Li, C.Y., and Yang, X.M. (2010b). PCBP-1 regulates alternative splicing of the CD44 gene and inhibits invasion in human hepatoma cell line HepG2 cells. Mol Cancer 9, 72.

Zhang, X.N., and Mount, S.M. (2009). Two alternatively spliced isoforms of the Arabidopsis SR45 protein have distinct roles during normal plant development. Plant Physiol 150, 1450–1458.

Zhang, Y., Xu, S., Ding, P., Wang, D., Cheng, Y.T., He, J., Gao, M., Xu, F., Li, Y., Zhu, Z., Li, X., and Zhang, Y. (2010c). Control of salicylic acid synthesis and systemic acquired resistance by two members of a plant-specific family of transcription factors. Proc Natl Acad Sci U S A 107, 18220–18225.

Zheng, X.Y., Spivey, N.W., Zeng, W., Liu, P.P., Fu, Z.Q., Klessig, D.F., He, S.Y., and Dong, X. (2012). Coronatine promotes Pseudomonas syringae virulence in plants by activating a signaling cascade that inhibits salicylic acid accumulation. Cell Host Microbe 11, 587–596.

Zipfel, C., Robatzek, S., Navarro, L., Oakeley, E.J., Jones, J.D., Felix, G., and Boller, T. (2004). Bacterial disease resistance in Arabidopsis through flagellin perception. Nature 428, 764–767.

